# Patterns of item nonresponse behavior to survey questionnaires are systematic and have a genetic basis

**DOI:** 10.1101/2022.02.11.480140

**Authors:** Gianmarco Mignogna, Caitlin E. Carey, Robbee Wedow, Nikolas Baya, Mattia Cordioli, Nicola Pirastu, Rino Bellocco, Michel G. Nivard, Benjamin M. Neale, Raymond K. Walters, Andrea Ganna

## Abstract

Response to survey questionnaires is vital for social and behavioral research, and most analyses assume full and accurate response by survey participants. However, nonresponse is common and impedes proper interpretation and generalizability of results. We examined item nonresponse behavior across 109 questionnaire items from the UK Biobank (UKB) (N=360,628). Phenotypic factor scores for two participant-selected nonresponse answers, “Prefer not to answer” (PNA) and “I don’t know” (IDK), each predicted participant nonresponse in follow-up surveys, controlling for education and self-reported general health. We performed genome-wide association studies on these factors and identified 39 genome-wide significant loci, and further validated these effects with polygenic scores in an independent study (N=3,414), gaining information that we could not have had from phenotypic data alone. PNA and IDK were highly genetically correlated with one another and with education, health, and income, although unique genetic effects were also observed for both PNA and IDK. We discuss how these effects may bias studies of traits correlated with nonresponse and how genetic analyses can further enhance our understanding of nonresponse behaviors in survey research, for instance by helping to correct for nonresponse bias.

## INTRODUCTION

Item nonresponse occurs when no substantive answer is recorded for a study participant on a given questionnaire item, such as when the participant doesn’t provide an answer or responds “I do not know”^1^. Nonresponse is interesting both as a behavioral choice by a survey participant and as a statistical concern due to missing data. Much social and behavioral research relies on surveys, and data analysis of survey data usually assumes full and accurate response by survey participants, or at least that any nonresponse is independent of the outcomes that a researcher is interested in. In reality, nonresponse is common and the independence assumption often unjustified, impeding proper interpretation and generalizability of results. Therefore understanding the causes of nonresponse has long been a concern for survey-based research^2^. The current study aims to evaluate item nonresponse patterns in a large prospective epidemiological cohort (the UK Biobank, or UKB) and clarify the contribution of genetics to differences in item nonresponse behaviors between individuals.

As an observable behavior, nonresponse represents a complex interplay between survey design for questionnaires and a respondent’s cognitive processes, i.e. in understanding a question and choosing a response^1,3^. Nonresponse at the item level may be thought of as an intermediate behavior on the spectrum between providing complete data and complete nonparticipation, i.e. unit nonresponse^4^, and nonresponse is predictive of future study dropout^5^. Further, nonresponse is unlikely to be captured by a single construct, since individuals may differ in their likelihood to select different nonresponse choices in a questionnaire, e.g. “I don’t know”, “I’m not sure”, or “I don’t want to answer”, both overall and when responding to questions in certain categories^6^.

Nonresponse behavior is also related to many other heritable traits. For example, higher rates of item nonresponse are correlated with lower educational attainment and poorer health status^7–9^. Increased item nonresponse has also been observed for individuals with more depressive symptoms^10^ and lower self-confidence, among other psychological and personality traits^11^.

However, these individual differences in item nonresponse rates can be sensitive to the content of the questionnaire items^7,12^ and the characteristics of the study population^9,13^. Still, similar patterns are often observed for unit nonresponse and study attrition, e.g., participants with lower educational levels are more likely to drop out of a study^14^. Similarly, those with heavy alcohol consumption or higher levels of mental distress tend to be underrepresented in studies due to their higher attrition rates^15–17^.

Understanding item nonresponse may address concerns about the generalizability of statistical analyses of observed data by helping to understand bias due to missingness. Item nonresponse reduces the effective sample size and can introduce bias^18^. Broadly, data may be either missing completely at random (MCAR), missing randomly conditional on the remaining observed data (missing at random, or MAR), or missing dependent on unobserved data (missing not at random, or MNAR)^6,19^. Thus if there are unobserved individual differences influencing the likelihood of item nonresponse, then the resulting missingness is considered MNAR. Nonrandom missingness is of particular concern because common statistical methods like full information maximum likelihood estimation or multiple imputation are only sufficient to address MAR data. MNAR requires more direct modeling that includes assumptions about the type of missingness^20–22^. Therefore, identifying a genetic component of item nonresponse behavior may assist with modeling MNAR mechanisms in genotyped samples^23^.

While other studies have focused on the genetic underpinnings of sex-differential participation or participation in optional study components^17,24–26^, the genetics of item nonresponse remain largely unknown. We first explore the phenotypic structure of the nonresponse options provided by the UK Biobank in the initial cohort assessment, and we estimate latent factors for a person’s general propensity to respond to questionnaires with “Prefer not to answer” (PNA) or “I don’t know” (IDK). We then perform GWAS of these two factors, identifying significantly associated loci and modeling genetic correlations with other heritable traits. We validate these genetic findings through out-of-sample polygenic prediction of nonresponse behavior. Throughout this investigation we avoid any analysis that would violate any participant’s stated desire to avoid answering a question (**Box 1**). We anticipate that these findings will provide insight into genetic influences on the cognitive processes involved in item nonresponse and also provide a basis for evaluating the impact of nonresponse bias on GWAS of other traits and disorders.

### BOX 1

Participant consent is critical for the ethical conduct of research. Nonresponse, including at the item level, in some instances will reflect a participant exercising their (entirely justified) right to voluntarily not participate in some aspect of a study. This is especially true in the case of item nonresponse in the form of actively responding “prefer not to answer”. As a result, it requires careful ethical consideration to evaluate how to study nonresponse without breaching the participant’s consent as reflected in both the item-level nonresponse and the study-level informed consent and participation.

There can be ethical harms to ignoring the source of missing data in research. Consideration of missingness is necessary to identify the ways in which a study or a particular analysis may not be representative of the population, otherwise researchers risk the myriad impacts of uncritically producing biased or ungeneralizable results. Decades of social science research on mechanisms of missing data and its influence on research results reflect recognition of the imperative to wrestle with this challenge.

With these considerations in mind, this paper endeavors to characterize broad, group level trends in overall amounts of item nonresponse while intentionally avoiding exploring nonresponse behavior (especially with regards to the “prefer not to answer” option) to single questions. We make one exception, in an analysis of responding “I don’t know” to the question “During your childhood, how many times did you suffer painful sunburn?”, where we empirically check that our analysis is meeting the stated goal of avoiding revealing item-specific factors. We intentionally rely on the IDK response (i.e., item nonresponse that doesn’t imply a desire to avoid participation in the item) and use an item that is less socially sensitive to minimize ethical concerns while doing this check.

The factors generated for our analyses can be thought of as reflecting a general behavioral tendency for someone to choose not to respond to one or more survey items; they are not reflective of nonresponse to any single, specific item. Perhaps most crucially, no attempts were made to infer individual item-level values of nonresponders, nor would such attempts be fruitful given the low predictive power of overall nonresponse behavior at the individual-person, individual-item level.

We stress that these ethical considerations apply not only within this study, but in future applications and extensions of this work. We encourage our colleagues to continue to remain vigilant to the challenges surrounding genetics and voluntary nonresponse in all domains.

## RESULTS

### Distribution of item nonresponse across questions

To investigate item nonresponse bias we considered two possible answer choices across 109 questions from the UK Biobank touchscreen questionnaire: “Prefer not to answer” and “I don’t know” (PNA and IDK, respectively). The final study population included 360,628 unrelated participants of European genetic ancestry with available genetic data. The PNA option was available for all questions, while only 83 questions allowed the IDK option. Participants selected PNA less frequently (8.82% at least once) than IDK (67.02% at least once) (**Tab. 1**, **Suppl. Fig. 1**). For each question, on average, 0.16% of participants chose PNA and 2.17% chose IDK (**Suppl. Tab. 1, Suppl. Tab. 2**). Importantly, individuals could only select one nonresponse answer per question, so a response of IDK necessarily precluded a response of PNA, and vice versa. Females answered PNA more often than males (57.15% females vs. 42.85% males, P<5×10^-5^), while females were only slightly more likely to answer IDK than males (52.90% females vs. 47.10% males, P<5×10^-5^) (**Tab. 1**). Nonresponders had markedly lower educational attainment (18.73% of nonresponders had college or university degrees vs. 33.45% for responders for PNA; 29.41% of nonresponders had college or university degrees vs. 37.75% for responders for IDK). (**Tab. 1**). PNA was the more common response among questions capturing potential illegal behavior or social stigma (e.g., “How often do you drive faster than the speed limit on the motorway?” or “Does your partner or a close relative or friend complain about your snoring?”) (**Suppl. Fig. 2**, panel **a**). Unsurprisingly, participants selected IDK more frequently in questions about their distant past such as “Were you breastfed when you were a baby?” or “During your childhood, how many times did you suffer painful sunburn?” (**Suppl. Fig. 2**, panel **b**). We hypothesized that the frequency of PNA or IDK answers might increase as a function of the order in which the questions were asked because of fatigue experienced by the participant as time spent taking a survey increases. We fit a negative binomial regression to explore this hypothesis and found no evidence of a positive trend for IDK or PNA.

**Tab. 1.** Baseline demographics about PNA and IDK nonresponders. PNA and IDK columns refer to those participants who chose these options at least once throughout the questionnaire.

| Characteristic |  | NO PNA<br>(N=328,843) | PNA<br>(N=31,785) | NO IDK<br>(N=118,928) | IDK<br>(N=241,700) |
| --- | --- | --- | --- | --- | --- |
| Mean age (SD) – year |  | 56.64 (8.00) | 58.55 (7.84) | 55.49 (8.08) | 57.46 (7.89) |
| Age group - no. (%) | ≤51yr | 93,859 (28.54) | 6678 (21.01) | 40,217 (33.82) | 60,320 (24.96) |
|  | 51<yr≤58 | 77,607 (23.60) | 6517 (20.50) | 29,220 (24.57) | 54,904 (22.72) |
|  | 58<yr≤63 | 80,082 (24.35) | 8151 (25.64) | 26,638 (22.40) | 61,595 (25.48) |
|  | yr>63 | 77,295 (23.51) | 10439 (32.84) | 22,853 (19.22) | 64,881 (26.84) |
| Female sex - no. (%) |  | 175,701 (53.43) | 18,166 (57.15) | 66,000 (55.50) | 127,867 (52.90) |
| Participants in UK Regions - no. (%) | East Midlands | 23,476 (7.14) | 2419 (7.61) | 8,336 (7.01) | 17,559 (7.26) |
|  | London | 21,953 (6.68) | 2139 (6.73) | 8,340 (7.01) | 15,752 (6.52) |
|  | North East | 38,720 (11.77) | 3990 (12.55) | 13,859 (11.65) | 28,851 (11.94) |
|  | North West | 50,212 (15.27) | 5276 (16.60) | 17,009 (14.30) | 38,479 (15.92) |
|  | Scotland | 25,216 (7.67) | 2416 (7.60) | 9,293 (7.81) | 18,339 (7.59) |
|  | South East | 30,905 (9.39) | 2603 (8.19) | 11,762 (9.89) | 21,746 (9.00) |
|  | South West | 45,626 (13.87) | 3659 (11.51) | 17,246 (14.50) | 32,039 (13.26) |
|  | Wales | 13,804 (4.20) | 1311 (4.12) | 4,981 (4.19) | 10,134 (4.19) |
|  | West Midlands | 28,499 (8.67) | 3186 (10.02) | 10,051 (8.45) | 21,634 (8.95) |
|  | Yorkshire | 50,432 (15.34) | 4786 (15.06) | 18,051 (15.18) | 37,167 (15.38) |
| College/University degree - no. (%) |  | 110,011 (33.45) | 5,924 (18.73) | 44,890 (37.75) | 71 ,075 (29.41) |

### Correlation patterns of item nonresponse and factor analyses

We used phenotypic (tetrachoric) correlations to measure the degree to which item nonresponse behavior is shared across survey questions (**Fig. 1**). PNA answers showed an overall higher correlation than IDK (mean r^2^=0.66 [interquartile range, or IQR=0.17] for PNA and mean r^2^=0.28 [IQR=0.13] for IDK), indicating that individuals who responded to questions with PNA tended to do so more consistently across questions that individuals who responded to questions with IDK. Indeed, we identified a small number of individuals who responded to all survey questions with PNA (N=11).

**Fig. 1.**
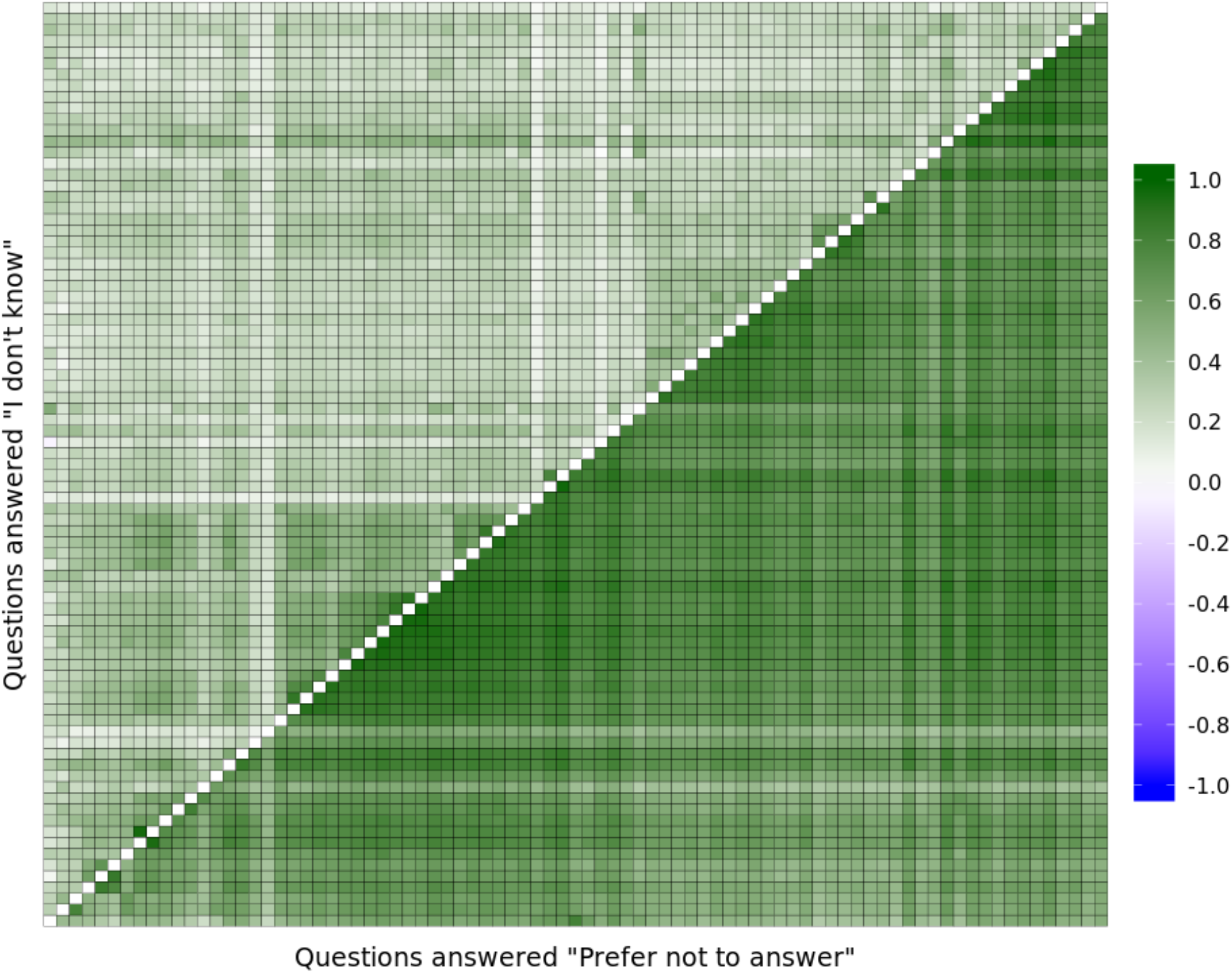
Phenotypic (tetrachoric) correlation of item nonresponse among questions. Each question has been recoded as dichotomous (1 = PNA or IDK, 0 otherwise). Each row and column represent the same question. We considered only questions which allow both the “Prefer not to answer” and the “I don’t know” options. The upper triangle = “I don’t know”, the lower triangle = “prefer not to answer”.

Item nonresponse behavior was also more similar among survey questions from similar phenotype domains. In other words, item nonresponse behavior was not independent of answering patterns across questions. For example, the average correlation of PNA answers among questions *within* the food intake and the mental health domains (mean r^2^=0.85 [IQR=0.08], and mean r^2^=0.76 [IQR=0.12], respectively) was higher than the correlations *between* food intake and mental health questions (mean r^2^=0.14 [IQR=0.05]).

Based on the observed structure across survey domains, we next estimated latent (unmeasured) factors for overall IDK and PNA item nonresponse behavior, respectively, conditional on the correlated substructure. To do so, for each response type we first assessed the survey substructure by performing factor analysis (FA) with the full set of questions and examining cluster analyses of the residual correlations from a single factor model (**Suppl. Fig. 3**, **Suppl. Fig. 4**, respectively). These residuals provided us with the magnitude of the correlation *not* explained by a single general factor model, allowing us to identify bi-factor FA model as an appropriate model for the survey substructure. The chosen bi-factor FA models the observed correlation matrix for item nonresponse as a function of one general factor affecting nonresponse for all items and possibly two or more additional domain-specific factors affecting subsets of items identified by the model. Since this model may not fully address nested substructure within groups of items we also evaluated fitting the bi-factor FA model on a reduced set of survey questions pruned for high pairwise correlations observed in the residual cluster analysis of the single factor model. From exploratory factor analysis we selected a 5 factor solution for the pruned PNA responses and a 4 factor solution for the pruned IDK responses - both with oblique (“biquartimin”) rotations – as our final models based on standard fit metrics (**Suppl. Tab. 3**).

We find that the common general latent factor, representing the underlying general item nonresponse behavior across questions, explained 51.26% and 25.61% of the total variance for PNA and IDK, respectively, based on the selected models. Our approach also identified substantial variance in item nonresponse behavior (11.63% and 11.20% for PNA and IDK, respectively) that was accounted for by additional domain-specific factors rather than a general factor (**Fig. 2**, for EFA: **Suppl. Fig. 6** and **Suppl. Fig. 7** for PNA and IDK, respectively). Two of these factors (influencing items we might think of as affecting “Health” and “Psychiatric” domains) partially overlap between PNA and IDK. The domain-specific factor with items related to “Ethnicity” was specific to PNA and was present when respondents did not answer question about ethnic background and skin color, with loadings of 0.69 and 0.51, respectively.

**Fig. 2.**
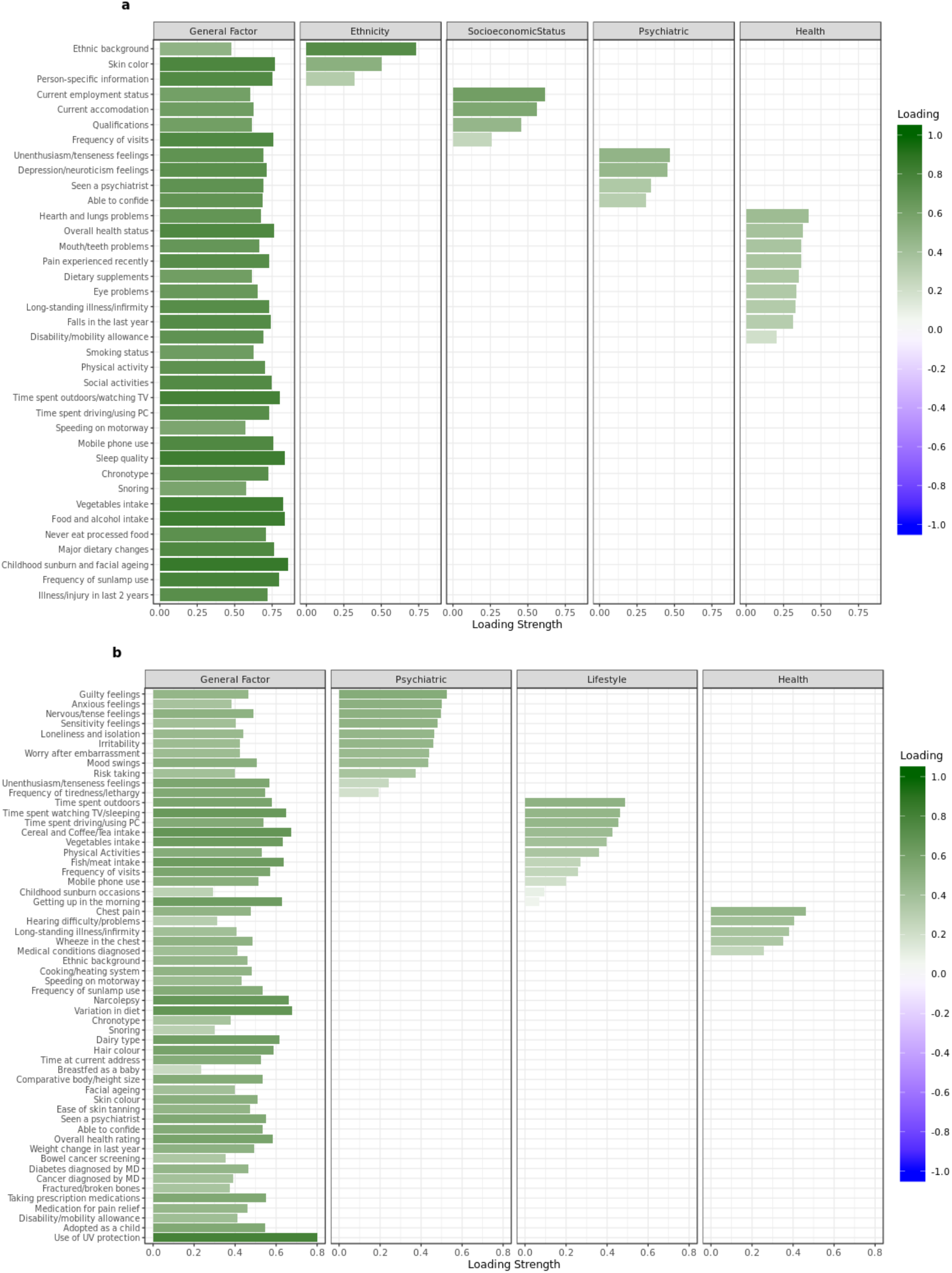
Bar graph of factor loadings of the questions in the PNA and IDK Confirmatory Factor Analyses. Panel **a** represents the loading strength of each question on the latent factors in the Exploratory Factor Analysis with a Bi-factor model for the “Prefer not to answer” analysis. Panel **b** represents the loading strength of each question on the latent factors in the Exploratory Factor Analysis with a Bi-factor model for the “I Don’t Know” analysis.

We then estimated each individual’s latent factor scores these general latent factors for PNA and IDK, respectively, in order to evaluate their relationship with other phenotypic and genetic measures. The density plots of the factor scores from the confirmatory factor analysis for both PNA and IDK can be seen in **Suppl. Fig. 5.**

### PNA and IDK factors predict response to follow-up questionnaires

We hypothesized that the common latent factors for PNA and IDK might capture a broader nonparticipation tendency beyond simply not answering specific survey questions. Therefore, we evaluated if the PNA and IDK latent factors were able to predict whether participants did or did not complete the online follow-up dietary questionnaires distributed by UK Biobank. We considered separately the completion of the follow-up online questionnaire only in the first wave of the invitation (N=69,735 and N=146,712 for PNA and IDK analyses, respectively) or in all the 4 waves of the invitation (N=19,097 and N=99,151 for PNA and IDK analyses, respectively). We quantified the improved prediction after controlling for age, sex, the top 20 principal components of the variance-covariance matrix of the genetic data, as well as education and self-rated health status since these have been shown to be proxies for survey nonresponse behaviors^14,24^. PNA and IDK factors slightly improved prediction of participation for all four waves of survey invitation, on top of education and self-rated health status, both when each factor was considered independently (incremental psuedo-R^2^=0.0027, P<2×10^-16^ and incremental pseudo-R^2^=0.0012, P=3×10^-11^, respectively for PNA and IDK), and when the two factors were combined (incremental pseudo-R^2^=0.0034, P<2×10^-16^). Combing PNA and IDK resulted in a better prediction of not responding to all four waves of follow-up survey invitations compared to predicting just one wave (incremental pseudo-R^2^=0.0012, P<2×10^-16^) (**Suppl. Tab. 4**). In summary, the general factors for item nonresponse are associated with whether participants will continue to engage in future, follow-up research, and our estimated scores for those factors are able to provide prediction beyond established proxies for nonresponse like education and self-rated health status.

### Genome-wide association study (GWAS) of item nonresponse

To assess potential genetic components of item nonresponse behavior we conducted a GWAS on the estimated factor scores for the general PNA and IDK behavior across survey questions. We identified 4 genome-wide significant (P<5×10^-8^) loci for PNA and 35 loci for IDK. (**Fig. 3** panel **a** and **b**, respectively).

**Fig. 3.**
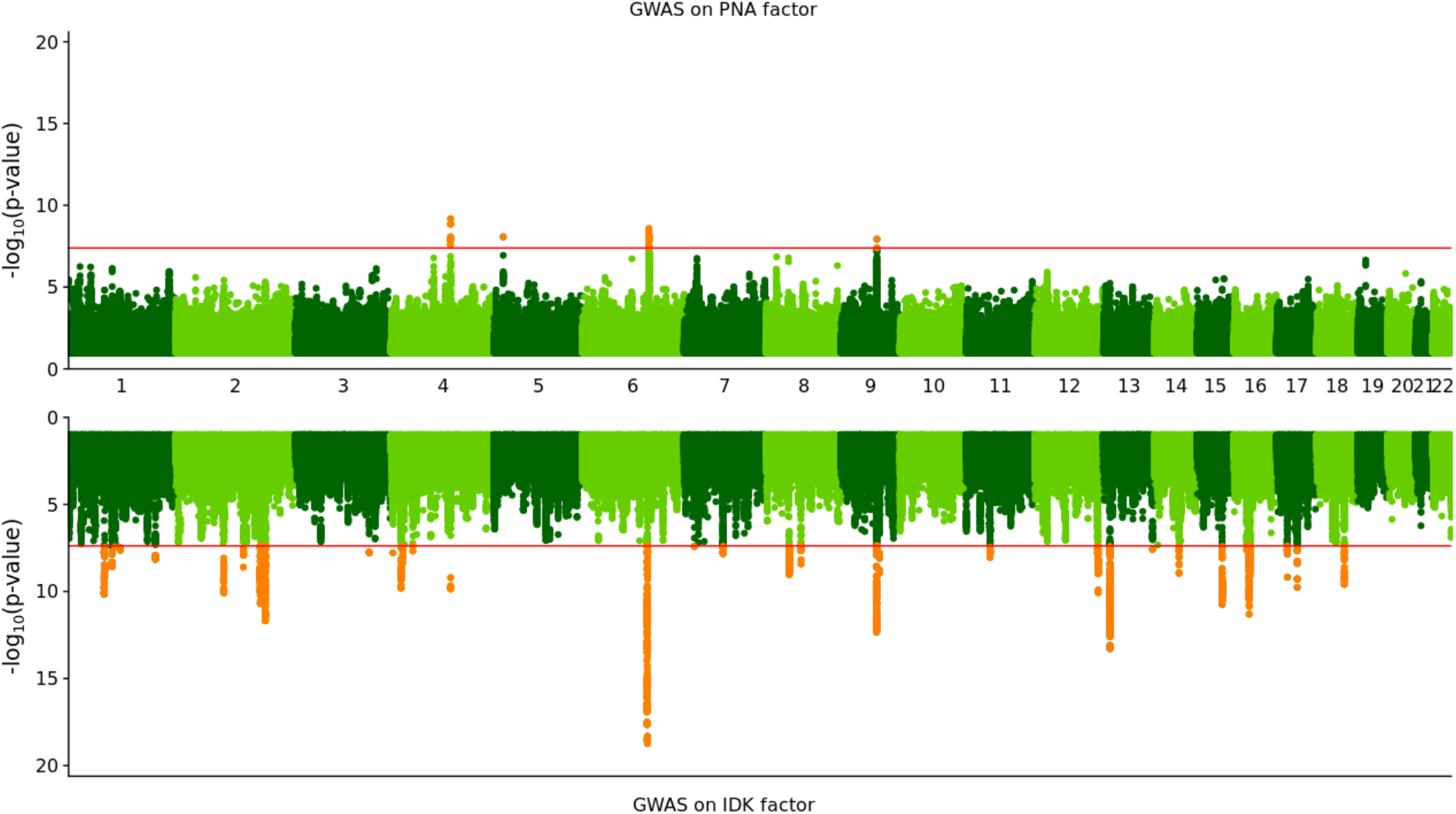
Miami Plot for GWAS of the “Prefer not to answer” and “I don’t know” factors. Miami plot of the resulting P-values from a GWAS of the PNA factor (on top) and the IDK factor (on bottom) for the 360,628 UK Biobank participants included in this study. The *x* axis is the chromosomal position, and the *y* axis represents the −log10 of the corresponding p-value for each location measured in the genome. The dashed line marks the threshold for genome-wide significance (*P* = 5×10^-8^).

Many of these genome-wide significant loci have been previously associated with traits related to poor health or lower socioeconomic status (SES). For example, allele T in rs79994730 is associated with an increased chance of answering PNA and IDK for survey questions, but also with lower educational attainment^27^ and intelligence^28,29^. Lead SNPs from 4 independent genomic loci overlapped with loci formerly associated with mental disorders^30,31^ and with gastrointestinal diseases^32,33^. For example, allele A in rs240764 is associated with more frequent neurotic behavior^30,32,34^. Similarly, allele A in rs13126505 is associated with an increased risk of developing Crohn’s disease^35^ and Inflammatory Bowel Disease^32^.

The general factors for PNA and IDK were both significantly heritable, with higher estimated SNP-heritability for IDK (*h^2^_g_*=.0675, P= 4×10^-112^) than for PNA (*h^2^_g_*=.0204, P=3×10^-16^). We also observed a significant heritability enrichment for brain tissue^36^ (**Suppl. Tab. 5, Suppl. Tab. 6**); the top two annotations were for Brain Cerebellum (P=3×10^-3^ for both PNA and IDK) and cerebellar hemisphere (P=5×10^-3^ and P=8×10^-3^ for PNA and IDK, respectively). Importantly, the SNP heritability for the factor scores shows substantially stronger genetic signal than GWAS using a simple sum of the number of nonresponses over all survey items. As comparison, the SNP heritability of the simple sum score for PNA responses was 13 times lower and nonsignificant (*h^2^_g_*=.0015, P=0.5). This is consistent with an expectation that the factor analysis provides improved power reducing measurement error across items and clarifying the signal in the context of correlated residual structure.

### Shared vs. question-specific item nonresponse behavior

One concern in our analysis is that the GWAS results for the item nonresponse phenotypes may be driven by questions with the highest number of PNA and IDK responses (**Suppl. Fig. 2**), rather than capturing an underlying behavior shared across survey questions. This is a concern both because it affects the interpretation of the results and because it could expose undesired information about nonresponse to individual items (**Box 1**). To explore this concern, we performed a GWAS of not knowing the answer to the question with the largest number of IDK responses, which was “During your childhood, how many times did you suffer painful sunburn?” We observed only a modest genetic correlation with the IDK factor (r_g_=0.39 [0.33,0.46]), and we identified 4 genome-wide significant loci for this GWAS. None of these 4 loci were genome-wide significant in the GWAS of either the PNA or IDK factors. As an example, rs35407-G allele, a 3 Prime UTR Variant in SLC45A2, is associated with a higher risk of *not* knowing how many times someone was sunburned as a child, and it also increases the risk for melanoma^37^ and cutaneous squamous cell carcinoma^38^. This result suggests that our factor score GWAS successfully highlights shared components affecting item nonresponse generally (*r_g_* > 0) while avoiding capturing more question-specific nonresponse behavior that is less related to overall nonresponse and therefore not the focus of our analyses.

### Genetic correlations with heritable traits

To better understand which traits and behaviors share genetic overlap with item nonresponse behavior, we calculated genetic correlations between the PNA and IDK factors with 655 different heritable phenotypes using LD score regression (**Fig. 4** and **Fig. 5** for PNA and IDK, respectively and **Suppl. Tab. 7** and **Suppl. Tab. 8** for PNA and IDK, respectively).

**Fig. 4.**
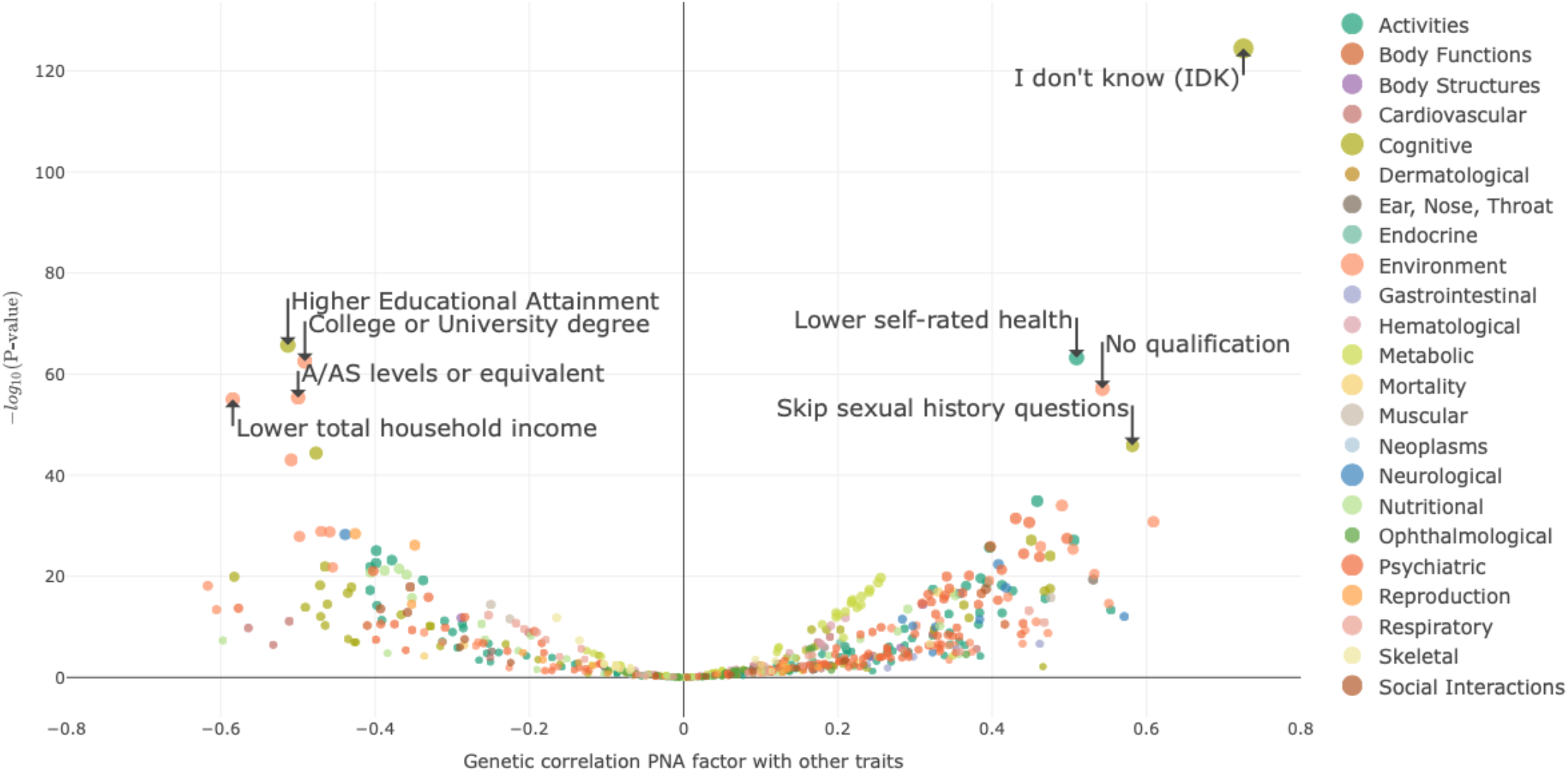
Volcano plots of the genetic correlation between the PNA factor and other heritable traits. The values on the x-axis represent the genetic correlation between the “Prefer Not to Answer” factor and 655 other heritable traits. The values on the y-axis represent the −log10 of the p-value of the associated statistical test. Only traits with a genetic correlation (in absolute value) greater than 0.45 and with P<10^-45^ are labeled.

**Fig. 5.**
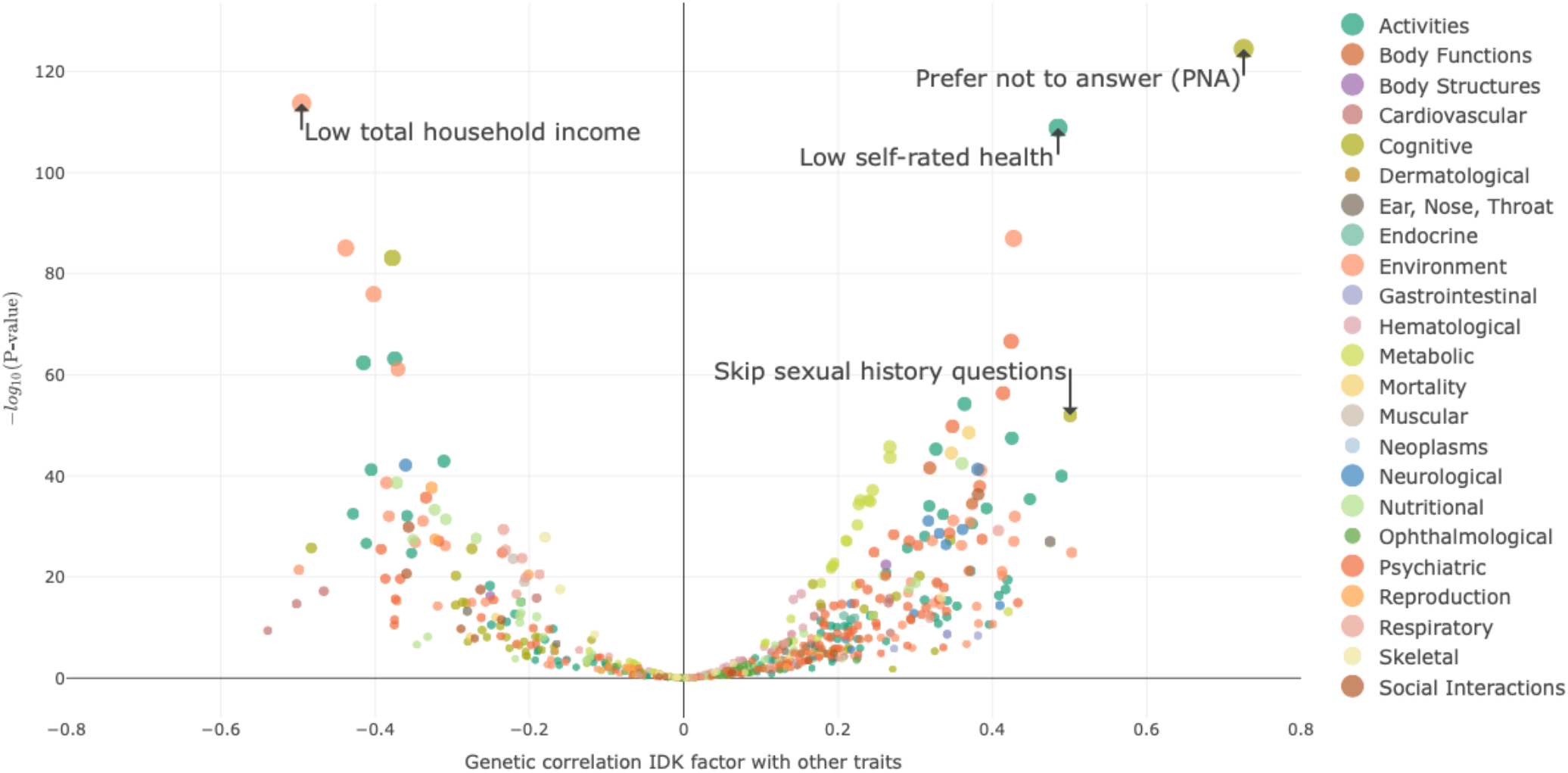
Volcano plots of the genetic correlation between the IDK factor and other heritable traits. The values on the x-axis represent the genetic correlation between the “I Don’t Know” factor and 655 other heritable traits. The values on the y-axis represent the −log10 of the p-value of the associated statistical test. Only traits with a genetic correlation (in absolute value) greater than 0.45 and with P<10^-45^ are labeled.

Overall, we observed a positive genetic correlation between the PNA and IDK factors and psychiatric disorders, poorer general health, lower educational attainment, and lower total household income before tax. These results are consistent with work from Adams and colleagues^14^ suggesting that nonresponse behaviors are strongly linked with socioeconomic status (SES). We highlight that only 109 out of 655 (16.64%) traits tested were included in the set of questions used to derive the item nonresponse phenotypes. For example, PNA and IDK factors had a high genetic correlation with opting to skip the sexual history section in the UKB survey (r_g_=0.58, [0.50,0.66], rg=0.50 [0.44,0.56] for PNA and IDK, respectively). This question was not used for deriving PNA and IDK factors, and it provides orthogonal evidence that the derived phenotypes are indeed capturing item nonresponse behavior in the UK Biobank.

We evaluated the degree to which the overall pattern of genetic correlations was driven by SES, we using Genomic Structural Equation Modeling^39,40^ (Genomic SEM). First, we calculated residual heritability of PNA and IDK conditional on genetic effects for total household income and educational attainment, since these are major elements of SES^41,42^. We estimate that resulting genetic variance unique to PNA and IDK was 1.33% (P=2×10^-16^) of the phenotypic variance in PNA and 5.09% (P=9×10^-78^) of the variance in IDK is attributable to unique genetic factors not explained by household income and educational attainment, corresponding to 65% and 75% of the total SNP heritability for PNA and IDK, respectively. We then estimated genetic correlations between PNA, IDK, and the remaining 654 heritable phenotypes with the same control for income and educational attainment with Genomic SEM. Overall, we observed a decrease in the number of traits significantly correlated with PNA and IDK factors after performing this analysis (**Suppl. Fig. 8**, **Suppl. Tab. 7**, **Suppl. Tab. 8**). However, both the PNA and IDK factors remained associated with poor self-reported health (r_g_=0.22 [0.14,0.30] and r_g_=0.27 [0.22,0.32], respectively) (**Suppl. Fig. 8**, **Suppl. Fig. 8**, **Suppl. Tab. 7**, **Suppl. Tab. 8**). SES-adjusted PNA also continued to be associated with the UKB measure for “having seen a psychiatrist for mental disorders” (r_g_=0.23 [0.15,0.31]). These results suggest that even after accounting for the genetic associations with socioeconomic factors, genetic associates with nonresponse were shared with genetic associations for poor overall health. Overall, these results highlight the utility of studying the genetics of item nonresponse, since these analyses help to assess which questionnaire items are likely to be affected by bias from nonrandom missingness, beyond what we can learn from using phenotypic information alone.

### Independent effect of PNA and IDK

In addition to the genetic correlation of the PNA and IDK factors with other traits, we observe substantial genetic correlation between these two factors (r_g_=0.73, [0.70, 0.76]), reflecting partial - but not complete (r_g_ <1) - genetic overlap between these two factors. Notably, genetic correlation facilitates this comparison between forms of item nonresponse by using genetics to overcome the limitation that UK Biobank participants could only respond with one of PNA or IDK on a given item, not both.

To help understand which genetic signals are unique to PNA and IDK, we estimated the conditional genetic correlation between PNA and other heritable phenotypes controlling for the genetic effects of IDK, and vice versa for correlations with IDK conditional on PNA, using Genomic SEM^40^. After accounting for the shared genetic associations between PNA and IDK, much of the genetic correlation observed between IDK and SES-related traits was reduced. Conversely, after accounting for the shared genetic associations between PNA and IDK, PNA preserved a more independent effect (**Suppl. Fig. 9**). The *PNA-adjusted* IDK factor was no longer associated with lower educational attainment (EA, r_g_=-0.01 [-0.07,0.05]) and the association with lower total household income (r_g_=-0.13 [-0.20,-0.06]) and poorer self-rated general health (r_g_=0.20 [0.13,0.27]) was attenuated. Conversely, *IDK-adjusted* PNA maintained a significant association with lower EA (r_g_=-0.37 [-0.31,-0.43]), low income (r_g_=-0.37 [-0.31,-0.43]) and poorer general health (r_g_=0.26 [0.21,0.31]). Moreover, *IDK-adjusted* PNA became associated with bipolar disorder (r_g_=0.25 [0.13,0.37]) and schizophrenia (r_g_=0.30 [0.23,0.37]). These results highlight that the genetic associations between PNA and IDK partially overlap, for instance in effects that are correlated with SES-related measures, suggesting shared influences across forms of item nonresponse. These results also highlight that the genetic associations between PNA and IDK are partially distinct, with PNA showing a unique overlap with psychiatric diseases that is not observed for IDK.

### Polygenic risk score analysis in Add Health

Finally, in order to test the generalizability of our genetic findings we constructed polygenic scores for the PNA and IDK factors in Wave 4 of the National Longitudinal Study of Adolescent to Adult Health (Add Health) data. Item nonresponse in Add Health was identified based on 163 questions with a possible response of “I don’t know” and 217 questions with a possible response of “refused to answer” (**Suppl. Tab. 9**). The IDK and PNA polygenic scores showed significant association with whether individuals a corresponding IDK or PNA response to at least one question. Specifically, we estimate that a one-standard-deviation increase in the PNA polygenic score is associated with a 2% increase in the probability of an individual ever answering with “refused to answer” in the Wave 4 Add Health data (incremental pseudo-R^2^ = 0.1%). We also estimate that a one-standard-deviation increase in the IDK polygenic score is associated with a 2% increase in the probability of an individual ever answering with “I don’t know” in the Wave 4 Add Health data (incremental pseudo-R^2^ = 0.5%).

We used a recently developed polygenic score for educational attainment (continuous years of completed education)^27^ derived from a very large sample size (N=1,131,881) to predict our two nonresponse outcomes in Add Health (**Suppl. Tab. 10**). We found that the educational attainment polygenic score is significantly associated with a 1% increase in the probability of an individual ever answering “refused to answer” in the Wave 4 Add Health data. The associated incremental pseudo-R^2^ is 0.08%. However, the educational attainment polygenic score was not significantly associated with the “I don’t know” outcome in the Wave 4 Add Health data. Taken together, these results suggest that our findings in UK Biobank replicate in an external US-based study of younger individuals.

## DISCUSSION

Nonresponse can impact the generalizability and reliability of survey-based research^43,44^. We show that item-level nonresponse is not random in the UK Biobank. Study participants vary in their degree and type of nonresponse, with observed differences between response patterns for PNA and IDK nonresponse options; the latter is more commonly chosen by study participants. Conversely, PNA responses are less common, but individuals who select PNA on at least one question are more consistent in their nonresponse patterns across questions. In characterizing these patterns of item nonresponse it is critically important to respect the ethical boundaries presented by the individual’s stated decision not to respond to a given item (**Box 1**). The current analysis evaluates overall item nonresponse behavior to identify genetic associations that are informative about nonrandom missingness in UK Biobank without implicating item-specific reasons for nonresponse.

Item nonresponse shows distinct correlation across questions, both broadly and within clusters that reflect the survey’s content and structure. Based on that observation, we’ve successfully characterized a “general” factor for each response pattern (i.e., PNA and IDK) that describes a broader nonresponse tendency across survey questions, while accounting for the presence of additional domain-specific correlations. Consistent with prior observations that item nonresponse is correlated with educational attainment, socioeconomic status, and overall health of participants who initially participated in a study^25^ or who continue to participate in follow-up waves of a study^26^, our PNA and IDK factors were associated with reduced educational attainment and predictive of response to follow-up surveys.

We also identified a significant genetic signal underlying our general nonresponse factors. We were able to use these results to study broad, group-level characteristics of individuals that did not respond to questions in the UK Biobank. The genetic correlation between PNA and IDK was high, but not 1, indicating that there are both shared and unique genetic associations for PNA and IDK. By using genetic structural equation models, we showed that the *PNA-specific* genetic signal was positively associated with psychiatric disorders, while this was not the case for the *IDK-specific* signal. Future work should focus on identifying both shared and unique genetic signals across different measures of nonresponse behaviors. Whether these results from UK Biobank are generalizable to other studies remains a matter for future investigation. However, our polygenic prediction results in Add Health suggest that at least part of the genetic signal underlying item-level nonresponse might be shared with other studies, and some of the signal is independent of other outcomes like educational attainment.

While such characterization is traditionally done by examining phenotypic correlations^45^, genetic analyses of our nonresponse outcomes provide some additional advantages beyond phenotypic data. For example, these analyses allowed us to characterize what individuals may be underrepresented in the study results beyond the phenotypic information that was collected in the survey questionnaires. While national registries or other administrative sources can sometimes be informative about nonresponse, either by comparing study participants to expected population descriptive statistics or by linkage to individual registry data^46^, these resources are not always available to researchers. Our analyses, in contrast, allowed us to leverage genetic information to characterize behavioral patterns of nonresponse. For instance, we were able to measure genetic correlations between survey nonresponse and GWAS summary statistics for hundreds of traits. Perhaps more importantly, these genetic correlations may highlight phenotypes that are likely to be affected by nonresponse bias in analyses that have not considered potential nonrandom missingness. Many genetic studies simply exclude participants who do not respond to surveys. Future work should consider opportunities for leveraging genetic information about item nonresponse to reduce nonresponse bias in analyses at the group level^23^.

In conclusion, we use phenotypic and genetic data to provide an extensive investigation of overall item-level nonresponse across items in the UK Biobank. These results should be considered when analyzing the UK Biobank, among other biobank-scale survey efforts, and when developing novel methods aimed at correcting and leveraging nonresponse in genetic analyses.

## METHODS

### UK Biobank and inclusion criteria

The UK Biobank is a health resource which has the purpose of improving the prevention, diagnosis, and treatment of a wide range of illnesses. It consists of a prospective cohort of 502,620 men and women aged 40-69 recruited in the years 2006-2010 throughout the United Kingdom. The touchscreen questionnaire is a collection of self-reported information regarding general health, dietary habits, physical activity, psychological and cognitive states, sociodemographic factors, etc. We began our inclusion criteria with 361,194 unrelated individuals of European genetic ancestry who passed quality control measures^47^. We excluded individuals who were enrolled only in the UKB pilot study (N=335). Participants who decided to terminate the touch screen questionnaire were asked to select PNA to all subsequent questions, and they were kept in our analyses. Conversely, individuals who withdrew from the study without filling out the touchscreen survey excluded from the analysis (N=231). As a result, a total of N=360,628 participants took part in the survey and answered every question of interest in the study; this is the final analytic sample size.

### National Longitudinal Study of Adolescent to Adult Health (Add Health) cohort

Add Health originated as an in-school survey of a nationally representative sample of US adolescents enrolled in grades 7 through 12 during the 1994-1995 school year^48^. Respondents were born between 1974 and 1983, and a subset of the original Add Health respondents has been followed up with in-home interviews, which allows researchers to assess correlates of outcomes in the transition to early adulthood. In Add Health, the mean birth year of respondents is 1979 (SD = 1.8), and the mean age at the time of assessment (Wave 4) is 29.0 years (SD = 1.8). All phenotypes included in this study come from Wave 4, the latest wave of Add Health data collection (2007-2009).

### PNA/IDK definitions

We considered only the Touchscreen questionnaire phenotypes with the PNA and IDK options (−3 and −1, respectively). We first studied the questionnaire protocol (https://www.ukbiobank.ac.uk/wp-content/uploads/2019/09/Touchscreen-questionnaire-for-website_Copyright.pdf) and kept only those questions asked to every study participants (N questions=109 and N=83 for PNA and IDK, respectively.). Questions asked to a subset of participants conditional on their answer to other questions were excluded.

### Phenotype definitions in Add Health

To investigate item nonresponse bias phenotypes in Add Health we considered two possible answer choices across hundreds of questions from the Wave 4 Add Health In-Home Interview questionnaire: “Refused to answer” and “I don’t know”. The final study population included 3,414 unrelated participants of European ancestry with available genetic data. The “refused to answer” option was available for 217 questions while only 163 questions allowed the “I don’t know” option. Our final outcomes were whether respondents ever answered at least once with “refused to answer” or “I don’t know”, respectively. We also predicted the two nonresponse outcomes in Add Health using a recently developed polygenic score for educational attainment^27^ (completed years of education).

### Factor Model Construction

#### Single-Factor Model

Exploratory Factor Analysis (EFA) with a single latent factor was performed separately on tetrachoric correlation matrices between each of the dichotomized PNA and IDK responses, respectively, and was implemented using the *fa* function from the “psych” package in R software version 3.4.4 with the oblique rotation “biquartimin” and the “Ordinary Least Square” extraction method.

Residuals from the initial EFA revealed a correlation structure indicative of further clustering unaccounted for by the general factor, with both broad correlations across item domains as well as some highly specific pairwise structure at the item level. Given that we were interested in modelling overall nonresponse behavior, not behavior specific to or driven by single item groupings or domains, we sought to reduce this additional structure first through the pruning of items with highly-correlated nonresponse patterns and then through the fitting of a bi-factor model. To further reduce the correlation structure between individual pairs and groups of variables, we considered different exploratory cut points in the dendrogram constructed on residuals from the single-factor EFA (**Suppl. Fig. 3**, **Suppl. Fig. 4**). We cut the dendrograms at height 0.500 and 0.775 in PNA and IDK, respectively, in order to minimize the number of branches (i.e., clusters of variables grouped together), but also maintain homogeneity within these branches (e.g., questions belonging to the same field). This led to 37 and 56 branches in PNA and IDK, respectively. In the IDK analysis, summing the IDK for each participant across questions inside each branch was sufficient to reduce the number of questions. We applied the same thresholds for PNA, but we noticed that in the four largest branches (i.e., person-specific information, food intake, overall health status, and mental health) the distribution of PNA per participant was J-shaped, with a continuously decreasing number of participants who chose PNA in more than 1 question and a small “peak” number of individuals who chose PNA in *every* question in the branch. For this reason, we scored each participant as follows: “0” if a participant answered all questions, “1” if a participant chose only PNA only once, “3” if a participant preferred not to answer *all* the items that fell in the same node, and “2” for participants who did not fit into the previous three categories. These scores were ordinal values used as input for bi-factor analysis that allowed for minimal item nonresponse information loss.

### Bi-Factor Analysis

To run bi-factor analysis^49,50^ on the pruned set of UKB questions we first split the dataset between 80% of participants (N=288,502) for Exploratory Factor Analysis^51^ (EFA) and 20% of participants (N=72,126) for Confirmatory Factor Analysis (CFA). For EFA we used the *fa* function, with “biquartimin” and “OLS” as the rotation and factoring method, respectively. We implemented CFA using the *cfa* function from the “lavaan” package^52^ in R software version 3.4.4, and also using the weighted least square mean and variance adjusted (WLSMV) estimator. We selected the initial factor structure from the EFA, first fitting models with different number of domain factors (**Suppl. Tab. 3**), then confirmed the fit of the model in the hold-out sample using the Root Mean Square Error of Approximation (RMSEA) and Tucker-Lewis Index (TLI). Upon selecting the optimal model and confirming fit, we re-ran the CFA in the full combined dataset; the final PNA and IDK phenotypes used in all downstream analyses were obtained as factor scores of the CFA-derived general factor in the full dataset. We extracted the factor scores using the “Empirical Bayes Modal” method as implemented in the *lavPredict* function.

### Predicting participation in follow-up questionnaires

We ran logistic regression to predict completion of an online follow-up 24-hour recall dietary questionnaire (field 110001 in the UKB) by using our PNA and IDK factors as predictor variable. To measure the variance explained by the model we computed the Pseudo-R^2^ using the McKelvey & Zavoina statistical method^53^. Completion of the first wave of a dietary questionnaire was coded as 1 if a participant completed this wave, and 0 if a participant did not (N=69,735 and N=146,712, respectively). Missing values (NA) were removed from the analysis (N=144,181). Similarly, completion of all 4 waves of the dietary questionnaire was coded as 1 if someone completed all 4 wave, 0 if someone didn’t complete all 4 waves of the questionnaire (N=19,097 and N=99,151, respectively). Other participants were not considered in this analysis (N=242,380). We examined the association of our standardized factors with sex, age, age^2^, sex-x-age^2^, the first 20 principal components of the variance-covariance matrix of the genetic data, self-reported health (field 2178 in the UKB), years of education. The years of education was created by recoding the Qualifications field (field 6138 in UKB) as follows^27^:

1. College or University degree (ISCED) = 20 years of education
2. A levels/AS levels or equivalent (ISCED 3) = 15 years of education
3. O levels/GCSEs or equivalent (ISCED 2) = 13 years of education
4. CSEs or equivalent (ISCED 2) = 12 years of education
5. NVQ or HND or HNC or equivalent (ISCED 5) = 19 years of education
6. Other prof. qual. (e.g., nursing, teaching) (ISCED 4) = 17 years of education
7. None of the above (ISCED 1) = 6 years of education

Participants who chose PNA or IDK for either years of education or self-reported health were excluded.

### Genotyping and Imputation

Genotyping and imputation procedures for the UK Biobank are detailed in Bycroft et al. 2018^54^. Genotyping in Add Health was performed at the Institute for Behavioral Genetics in Boulder, CO, using Illumina’s Human Omni1-Quad-BeadChip^55^. After imputing the genetic data to the Haplotype Reference Consortium (HRC)^56^ using the Michigan Imputation Server^57^, only HapMap3 variants were included, which are well imputed and provide good coverage of common variation across the genome. Analyses were limited to individuals of European-ancestry, and cryptically related individuals and ancestry outliers were dropped from analyses. Finally, only HapMap3 variants with a call rate above 98% and a minor allele frequency > 1% were used.

### GWAS

We performed genome-wide association studies using a linear model implemented in Hail^58^, including sex, age, age^2^, sex-x-age^2^, and the first 20 principal components of the variance-covariance matrix of the genetic data. We included only variants with imputation INFO score > 0.8 and MAF > 0.01, resulting in N=1,089,173 total SNPs in our GWAS.

### Results from FUMA GWAS catalogue

We used the FUMA^59^ pipeline to identify independent genomic loci. We considered an independent locus as the region including all SNPs in pairwise Linkage Disequilibrium (r^2^ > 0.6), with the lead SNPs in a range of 250 kb and independent from other loci at r^2^ < 0.1. We used the 1000 Genomes Phase3 Northern Europeans LD reference panel.

### Heritability and tissue-specific heritability

We used GWAS summary statistics with LD Score regression^60^ to estimate the proportion of variation in a trait that is explained by inherited genetic single nucleotide polymorphisms (SNPs). The rationale behind this method is that, for a polygenic trait, the higher the Linkage Disequilibrium of a variant with other variants, the more likely the index variant will tag a causal variant, and therefore its resulting LD Score will be higher. We included the SNPs in the HapMap3 reference panel (N=1,217,312) as a reference set.

Stratified LD Score regression^61^ proceeds with the rational that the χ^2^ association test for an index SNP includes the effects of all the SNPs tagged by that index SNP. For a polygenic trait, the χ^2^ association test will be higher for SNPs in LD with the index SNP, which can occur either when SNPs in LD tag an individual large effect SNP or when they tag several weak SNP effects. By partitioning SNPs into functional categories, SNPs in LD with the index SNP will increase the χ^2^ association test more so than SNPs in LD with a given SNP that belongs to a different functional category.

### Genetic Correlation

The genetic correlation between traits using GWAS summary statistics was computed using LD Score regression^62^, using the same reference set of SNPs as was used to estimate heritability. Under a polygenic model, LD Score Regression posits that the GWAS effect size for each variant includes the effect of all the variants that the index variant tags. Therefore, the genetic covariance can be estimated using the slope from the regression of the product of the z-score of the same variant from different GWA studies. We ran genetic correlations for PNA and IDK with a total of 654 traits, 616 which were from the UK Biobank and publicly available^63^. Traits used for genetic correlation analyses were chosen before conducting the analyses, with the agreement of the coauthors.

### Genomic SEM

Genomic Structural Equation modeling (Genomic SEM)^39^ is a two-stage structural equal modeling approach. In the first stage, the genetic and sampling covariance matrices are estimated using the Diagonally Weighted Least Squares (DWLS) estimation procedure. In the second stage, a multivariate system of covariance associations involving the genetic components of phenotypes are specified, and their corresponding parameters are estimated by minimizing the discrepancy between the model-implied covariance matrix and the empirical covariance matrix. We used Genomic SEM to run genetic correlations between PNA and IDK and other traits that influence nonresponse, while adjusting for other correlated phenotypes^40^. In particular, we adjusted for educational attainment^27^, self-rated health^64^, and total household income before tax. Moreover, we computed genetic correlations with Genomic SEM between PNA and other traits, adjusting for IDK, and vice versa.

### Polygenic risk scoring

A polygenic score for an individual is a weighted sum of a person’s genotypes at *J* loci,

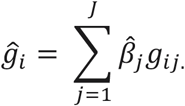

where 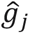 denotes the polygenic score of individual *i*, 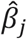 is the estimated additive effect size of the effect-coded allele at variant *j*, and *g_ij_* is the genotype of individual *i* at variant *j* (coded as having 0, 1, or 2 instances of the effect-coded allele). The polygenic scores were constructed with LDpred^65^, a method shown to have greater prediction accuracy than the conventional risk prediction approach involving LD pruning followed by *p-value* thresholding. LDpred considers the genetic architecture by accounting for LD among the SNPs in creating the polygenic scores. We used a Wald test to evaluate the significance of the polygenic scores on the outcomes.

For the Add Health sample, we used the genotyped data from the Add Health prediction cohort to create the LD reference file. After imputing the genetic data to the Haplotype Reference Consortium (HRC)^56^ using the Michigan Imputation Server^57^, we used only HapMap3 variants with a call rate > 98% and a minor allele frequency > 1% to construct the polygenic scores. We limited the analyses to European-ancestry individuals. Polygenic scores were calculated with an expected fraction of causal genetic markers set at 100%. In total, we used 1,168,025 HapMap3 variants to construct the polygenic scores in Add Health. We then used Plink^66^ to multiply the genotype probability of each variant by the corresponding LDpred posterior mean over all variants. In total, we created two polygenic risk scores, using the summary statistics of our two main phenotypes: 1) Prefer not to Answer (PNA) and 2) I Don’t Know (IDK). We then determined the association of the polygenic score for the related Refused to Answer and I Don’t Know Phenotypes in Add Health. Prediction accuracy was based on an ordinary least squares regression of the outcome phenotype on the polygenic score and a set of standard controls, which include birth year, sex, an interaction between birth year and sex, and the first 10 genetic principal components of the variance-covariance matrix of the genetic data. Variance explained by the polygenic risk scores was calculated in regression analyses as the R^2^ change (or Nagelkerke’s pseudo-R^2^ change for the dichotomous variables), i.e. the R^2^ of the model including polygenic risk scores and covariates minus the R^2^ of the model including only covariates. 95% confidence intervals around all R^2^ values are bootstrapped with 1000 repetitions each. We also used a recently developed score for educational attainment to predict both of our nonresponse outcomes in Add Health^25^.

## ACKNOWLEDGEMENTS

This research was conducted by using the UK Biobank Resource under application 31063. A.G. was supported by Academy of Finland (grant no. 323116) and by the European Research Council (ERC) under the European Union’s Horizon 2020 research and innovation programme (grant no. 945733). This project has also received funding from the European Union’s Horizon 2020 research and innovation programme under grant agreement no. 101016775. The National Longitudinal Study of Adolescent to Adult Health (Add Health) is supported by grant P01 HD031921 to Kathleen Mullan Harris from the Eunice Kennedy Shriver National Institute of Child Health and Human Development (NICHD), with cooperative funding from 23 other federal agencies and foundations. Add Health GWAS data were funded by NICHD grants to Harris (R01 HD073342) and to Harris, Boardman, and McQueen (R01 HD060726). For information about access to the data from this study, contact. We especially acknowledge and are grateful for the contributions of the participants in these studies, both for providing biological data and for their responses and nonresponses to survey questions that made this study possible.

## AUTHOR CONTRIBUTIONS

A.G. and R.K.W. designed and oversaw the study. G.M. was the study’s lead analyst, responsible for quality control, meta-analyses, and all major statistical analyses. Analysts who assisted G.M. in major ways include: C.E.C. (factor analysis), R.W. (prediction), N.B. (GWAS analyses), M.C. (statistical analyses), and R.B. (statistical analyses). N.P., M.G.N., and B.M.N. provided helpful advice and feedback on various aspects of the study design and analyses. All authors contributed to and critically reviewed the manuscript. G.M., C.E.C., R.W., R.K.W., and A.G. made especially major contributions to the writing and editing.

## COMPETING INTERESTS

Benjamin M. Neale is a member of the scientific advisory board at Deep Genomics and Neumora, consultant of the scientific advisory board for Camp4 Therapeutics and consultant for Merck.

## DATA AVAILABILITY

The GWAS results are available through the GWAS catalog accession nos. [GWAS SUMMARY STATISTICS ARE CURRENTLY BEING SUBMITTED TO THE GWAS CATALOGUE AND THIS SECTION WILL BE UPDATED WITH ACCESSION NUMBERS WHEN THEY ARE AVAILABLE.]

## CODE AVAILABILITY

All software used to perform these analyses are available online. Scripts used to perform these analyses are available at: https://github.com/gianmarcomigno/Item-nonresponse.

## Notes

https://github.com/gianmarcomigno/Item-nonresponse

