## Supplementary Figures for "Patterns of item nonresponse behavior to survey questionnaires are systematic and have a genetic basis"

**This PDF file includes:**

Supplementary Figures 1 to 9

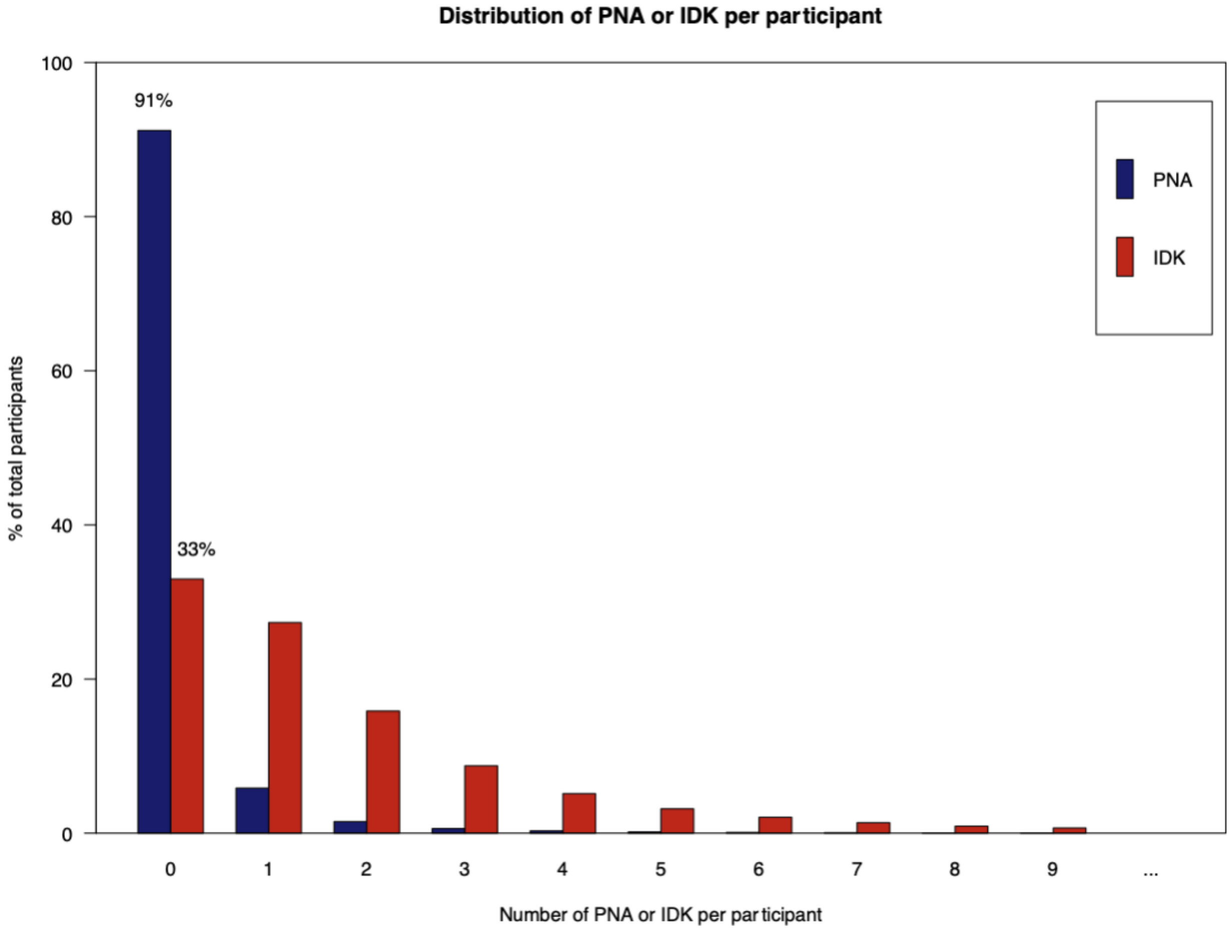


**Supplementary Figure 1. Distribution of PNA and IDK per participant in the full UKB cohort**

The plot highlights descriptives for different nonresponse behavior of participants in the PNA and IDK analyses. While 91% of the complete N=360,628 analytic sample reported no PNA throughout the UKB questionnaire, 33% reported no IDK throughout the UKB questionnaire. In other words, 9% of the sample preferred not to answer at least one question, whilst 67% of the same sample didn't know how to answer at least one question of the touchscreen questionnaire in UKB.

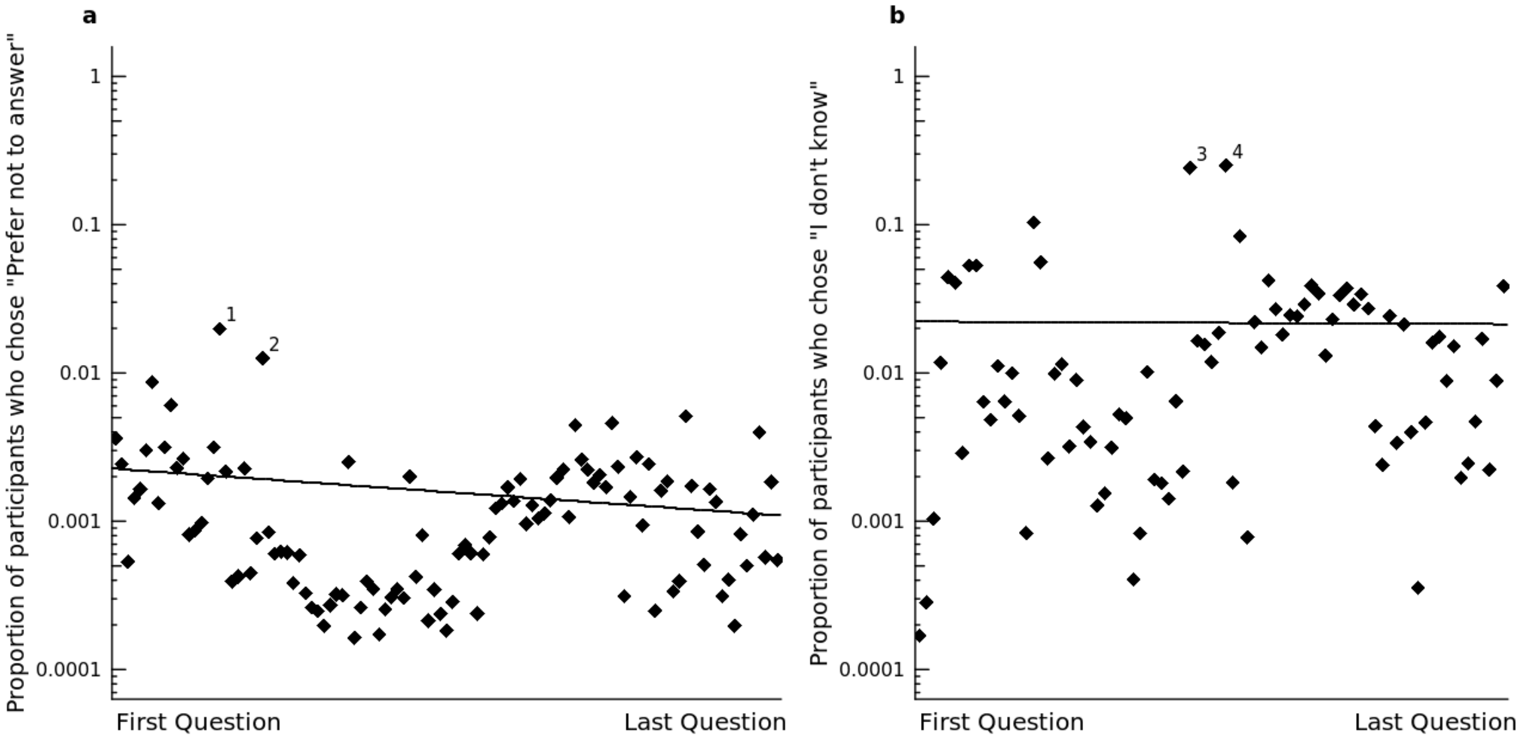


**Supplementary Figure 2. Distribution of item nonresponse across 109 UK Biobank touchscreen questionnaire questions**

Percentage of participants who chose the option "Prefer not to answer" (panel **a**) or "I don't know" (panel **b**). Questions have been ordered in the same order as they appeared in the questionnaire. The lines represent the fitted values from a negative binomial regression of the counts of PNA and IDK. Questions annotated are those with the highest number of nonresponses. **1**: "How often do you drive faster than the speed limit on the motorway?", **2**: "Does your partner or a close relative or friend complain about your snoring?", **3**: "Were you breastfed when you were a baby?", **4**: "Before the age of 15, how many times did you suffer sunburn that was painful for at least 2 days or caused blistering?"

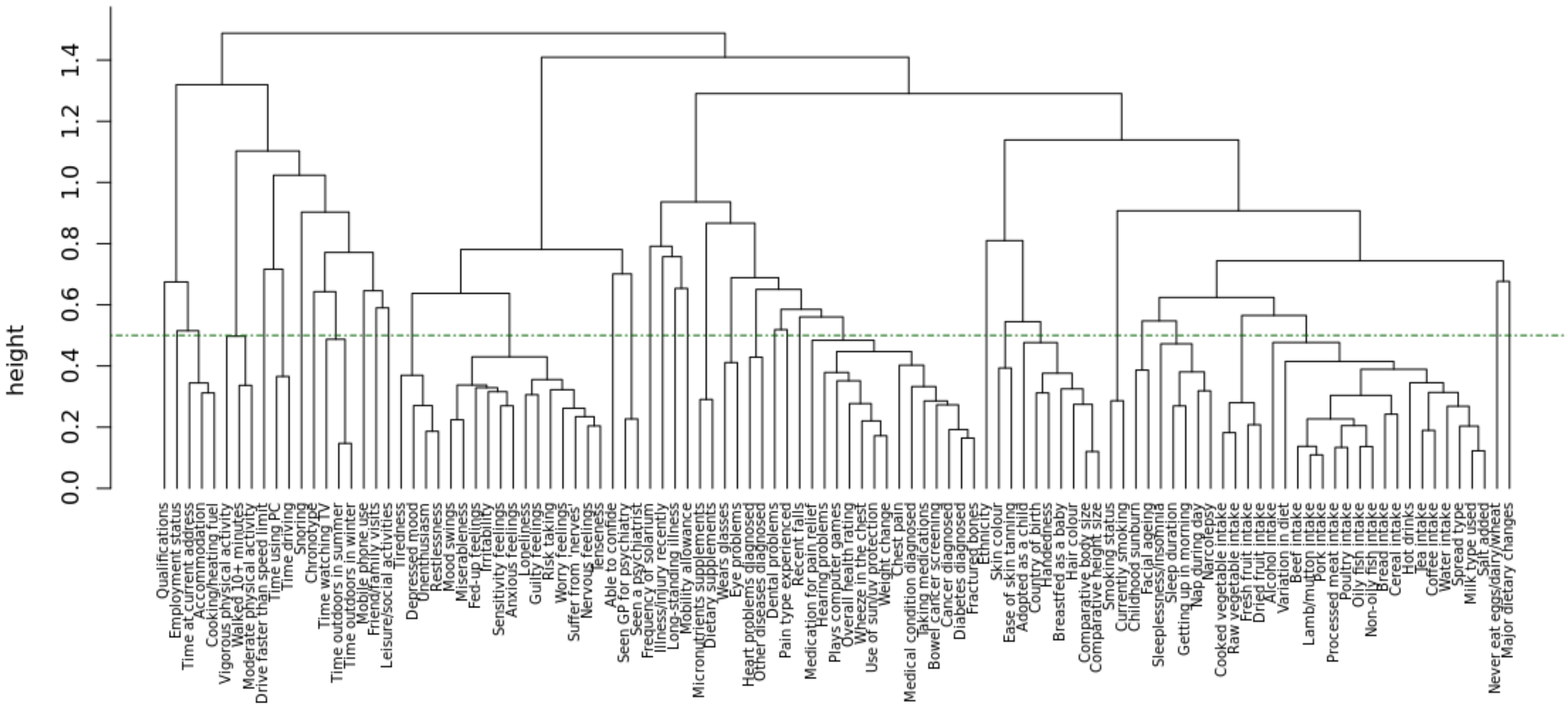


**Supplementary Figure 3. Dendrogram of residuals from Factor Analysis with one factor for PNA**

The dashed line at height 0.500 is the cut point that allowed us to reduce the number of questions to keep in the analysis.

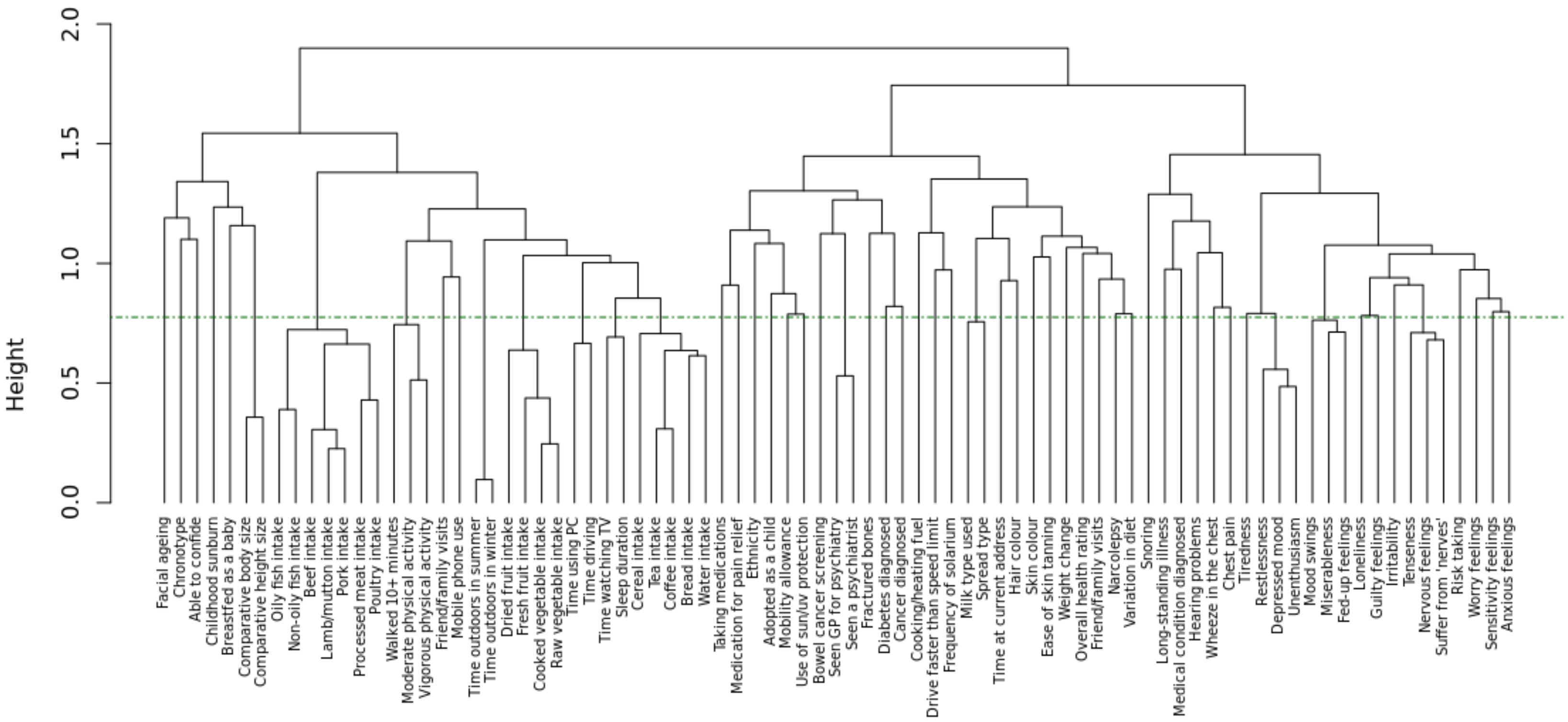


**Supplementary Figure 4. Dendrogram of residuals from Factor Analysis with one factor in for IDK**

The dashed line at height 0.775 is the cut point that allowed us to reduce the number of questions to keep in the analysis.

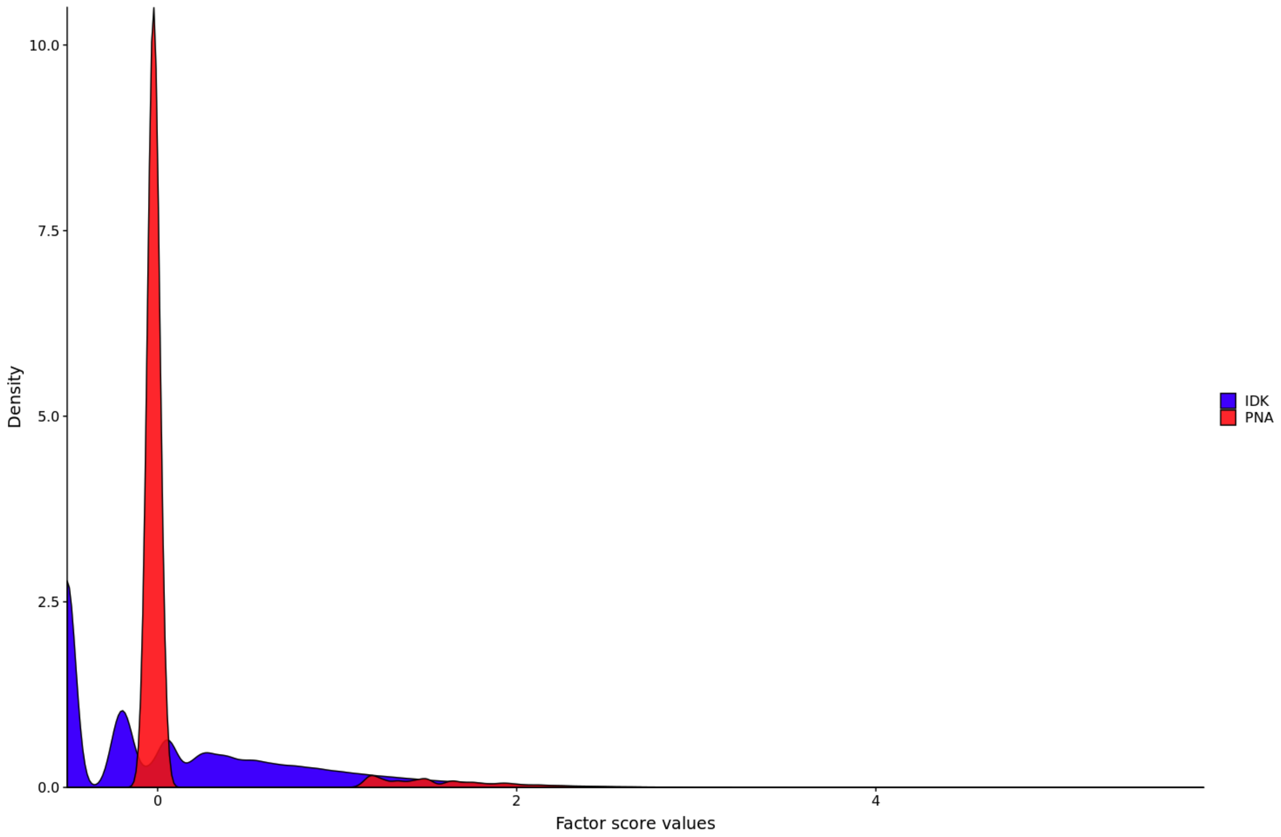


**Supplementary Figure 5. Density plots of factor scores from the Confirmatory Factor Analysis for both PNA and IDK.**

The plot shows the densities of the factor scores for the general latent factor from the Confirmatory Factor Analysis for both PNA and IDK in the touchscreen questionnaire in UKB. Factor scores for PNA mostly clustered around 0, while factor scores for IDK were more sparse, with less values clustered around 0.

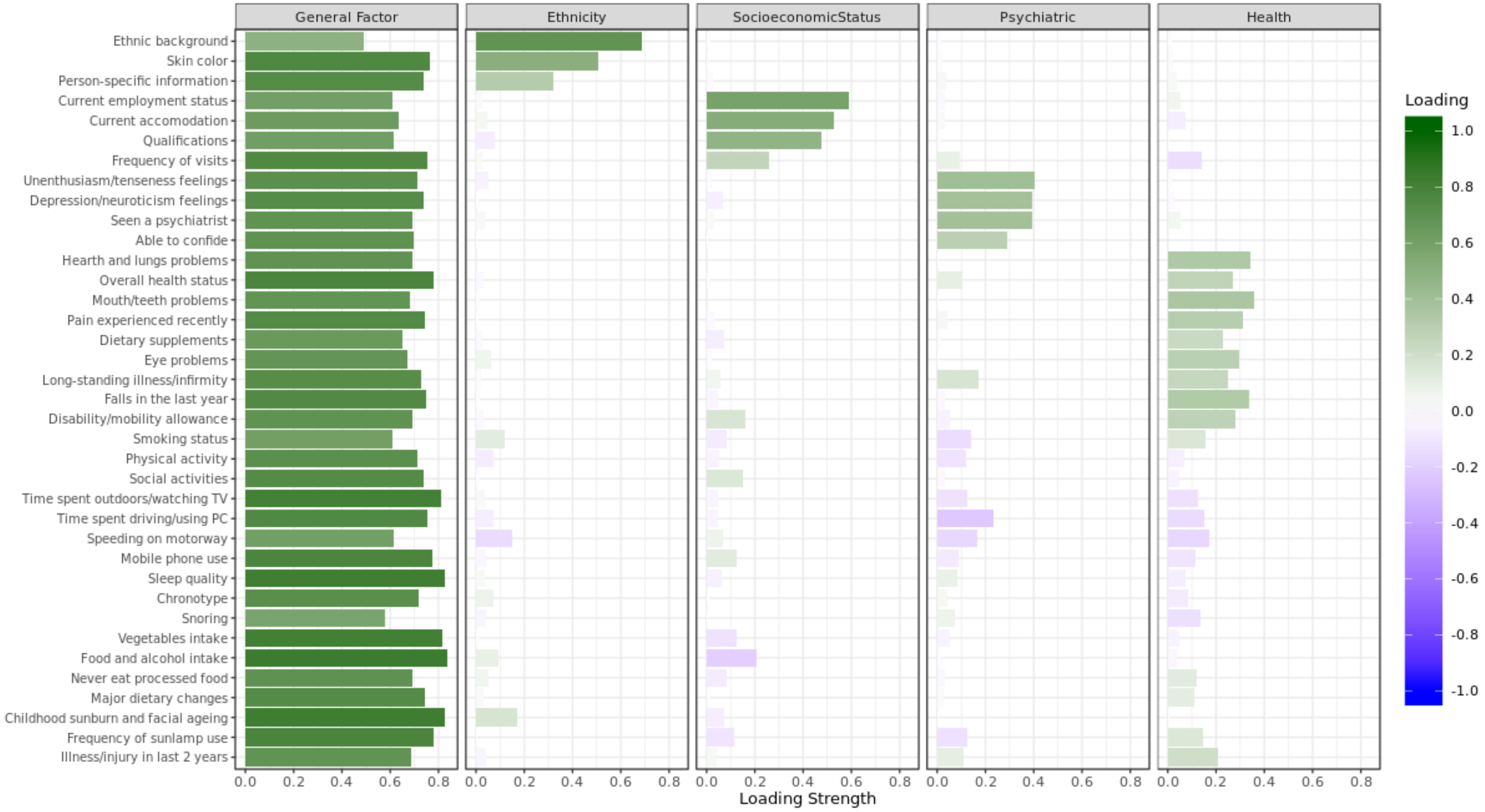


**Supplementary Figure 6. Bar graph of factor loadings for questions in the PNA Exploratory Factor Analysis**

The plot represents the loading strength of each question on the latent factors in the Exploratory Factor Analysis with a Bi-factor model in the "Prefer Not to Answer" analysis.

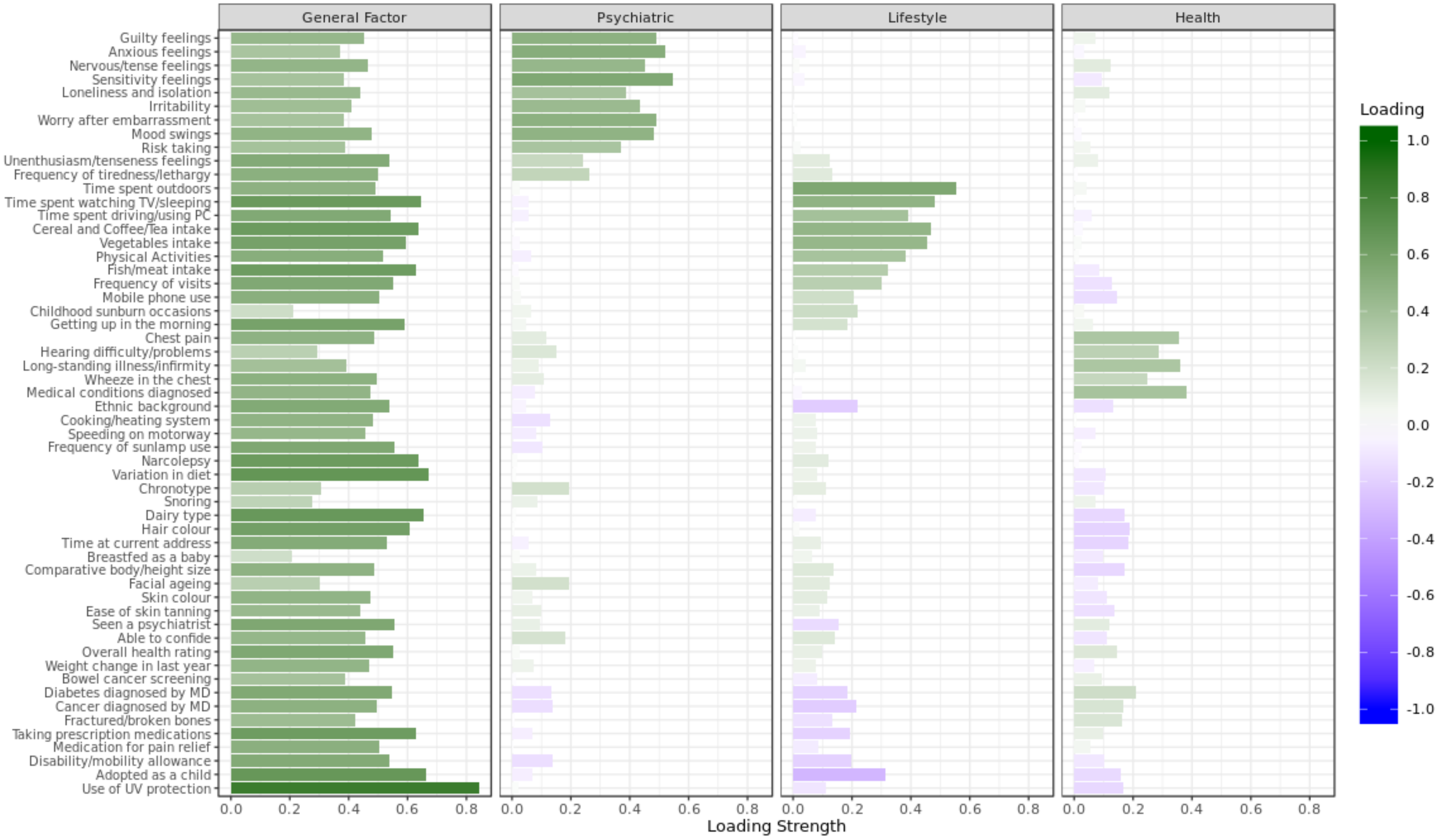


**Supplementary Figure 7. Bar graph of factor loadings for questions in the IDK Exploratory Factor Analysis**

The plot represents the loading strength of each question on the latent factors in the Exploratory Factor Analysis with a Bi-factor model in the "I Don't Know" analysis.

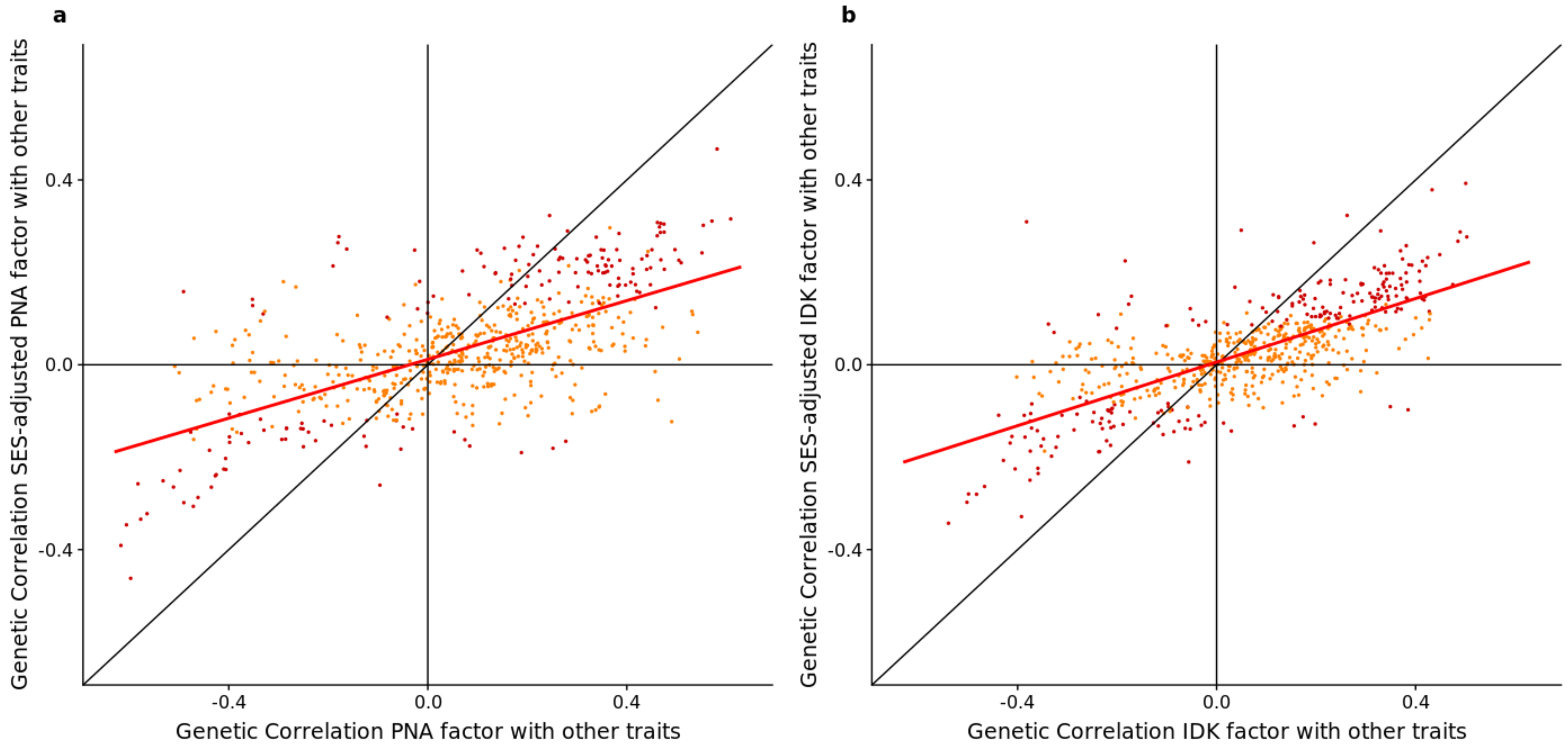


**Supplementary Figure 8. Adjusted vs unadjusted genetic correlations between PNA/IDK factors and other heritable traits, adjusted for SES.**

Comparison of the genetic correlation between PNA in panel **a** and IDK in panel **b** with other heritable traits, before and after (x and y axes, respectively) adjusting for Socioeconomic Status (SES), namely income and educational attainment, using Genomic SEM. Dots on the bisecting line suggest that the genetic correlation between PNA/IDK and other traits, after subtracting the effect of socioeconomic confounders, remains unchanged. Darker dots are those significant after a multiple hypothesis testing correction ($\alpha$=0.05, N traits=654). The red lines are regression lines, interpolating the dots.

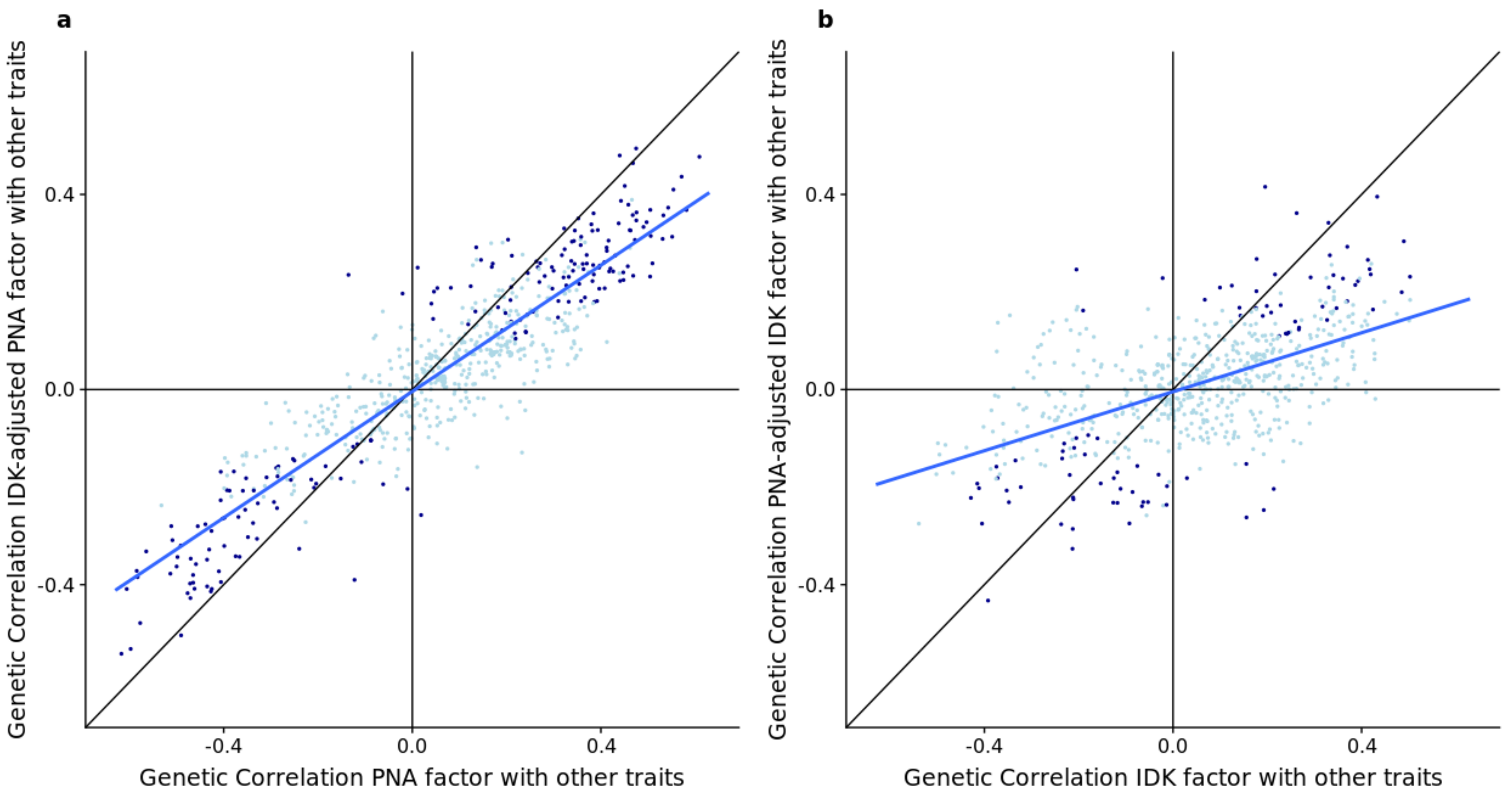


**Supplementary Figure 9. Adjusted vs unadjusted genetic correlations between PNA/IDK factors and other heritable traits, adjusted for IDK/PNA, respectively**

Comparison of the genetic correlation between PNA in panel **a** and IDK in panel **b** with other heritable traits, before and after (x and y axes, respectively) adjusting for PNA/IDK, respectively using Genomic SEM. By adjusting for PNA/IDK we remove the effect of PNA from IDK, and vice versa. Dots on the bisecting line suggest that the genetic correlation between PNA/IDK and other traits, after subtracting the effect of the other nonresponse trait, remains unchanged. Darker dots are those significant after a multiple hypothesis testing correction ($\alpha$ =0.05, N traits=654). The blue lines are regression lines, interpolating the dots. Since the line in panel **a** is closer to the bisector line, this suggests that most of the genetic correlations between PNA and the other traits remained unchanged, both before and after subtracting the effect of IDK.
